# The Big (Genetic) Sort? Reassessing Migration Patterns and Their Genetic Imprint in the UK

**DOI:** 10.1101/2023.02.27.530194

**Authors:** Shiro Furuya, Jihua Liu, Zhongxuan Sun, Qiongshi Lu, Jason M. Fletcher

**Affiliations:** Department of Sociology, University of Wisconsin–Madison, Madison, WI, 53706; Center for Demography of Health and Aging, University of Wisconsin–Madison, Madison, WI, 53706; Center for Demography and Ecology, University of Wisconsin–Madison, WI, 53706; Department of Biostatistics, Epidemiology, and Informatics, Perelman School of Medicine, University of Pennsylvania, PA, 19104; Department of Biostatistics and Medical Informatics, University of Wisconsin–Madison, WI, 53706; Department of Statistics, University of Wisconsin–Madison, Madison, WI, 53706; La Follette School of Public Affairs, University of Wisconsin–Madison, Madison, WI, 53706

## Abstract

This study reassesses Abdel Abdellaoui et al.’s findings that genetically selective migration may lead to persistent and accumulating socioeconomic and health inequalities between “types” (rich or poor) of places in the UK. Their work categorized migrants who moved to the same “type” of place (rich-to-rich or poor-to-poor) as non-migrants. We re-investigate the question of genetically selective migration by examining migration patterns between places rather than “place-types” and find genetic selectively in *whether* people migrate rather than *where*. For example, we find evidence of positive selection of people with genetic variants correlated better education moving from rich to poor places with our measure of migration that was obscured in the earlier work that used a non-standard measure of migration.

---

Geographic mobility (i.e., migration) is a process that shapes geographic distribution of a wide range of traits. Genetic predisposition is not an exception. Recent work by Abdel Abdellaoui et al. (2019) (henceforth, AA) in *Nature Human Behaviour* demonstrated the role of migration in geographic clustering of genetic variants, particularly alleles associated with educational attainment (EA). Specifically, those who moved to coal mining (i.e., economically deprived) places carry a low EA polygenic index (PGI), whereas those with a high EA PGI were selected to leave coal mining places. These migration patterns are consistent with the findings that skilled individuals are more likely migrate to seek better education and occupational opportunities (Coulter & Scott, 2015), while the durable and affordable housing and public transportation networks in coal mining places attract poor, low-skilled migrants (Rodden, 2010). Based on these results, AA provided two important implications: (1) own genetic variants are similar to their neighbors’ genetic variants than genetic variants of those living far away; and (2) these patterns of migration selection exacerbate geographic differences in genetic variants.

AA uncovered an important role of geographic mobility in genetic research on sociogeographic patterning; however, we point to a limitation in the operationalization of AA’s migration measure. AA’s analysis abstracted from typical measures of migration to only consider flows between “types” of places being either rich or poor; therefore, people who migrated between rich places or between poor places were classified as non-migrants. This departure from standard measures of migration blurs the interpretation of their analysis, as “non-migrants” are composed of migrants and non-migrants, and these latter two groups can be substantially dissimilar for a number of reasons demonstrated in previous work.

The healthy migrant hypothesis posits that skilled and healthy individuals are more likely selected to migrate than unskilled and less healthy counterparts (Jasso et al., 2004; Palloni & Arias, 2004; Palloni & Morenoff, 2006). Empirical evidence supporting the healthy migrant hypothesis has been repeatedly provided in the context of international (Crimmins et al., 2005; Fuller-Thomson et al., 2015; Landale et al., 2000; Riosmena et al., 2013; Rubalcava et al., 2008) and internal migration (Lu, 2008; Ma et al., 2020; Molloy et al., 2011; Nauman et al., 2015; Tong & Piotrowski, 2012; Wilding et al., 2016). Additionally, empirical evidence for the positive associations between migration distance and skills and health (Boyle et al., 2001; Liu & Shen, 2017; McCollum et al., 2021; Wilding et al., 2018) also suggests potential differences between migrants and non-migrants. These consistent findings for differences between migrants and non-migrants imply the importance of a clear distinction by whether people moved in migration research. Nonetheless, AA’s migration measure does not distinguish this set of migrants from non-migrants. This combined comparison category does not allow the authors to distinguish between genetic selection of whether to migrate versus where to migrate. The goal of the current study is to deepen our understanding of the relationship between migration, social stratification, and genetic measures through the reassessment of AA’s work. With our operationalization of migration, we found that non-migrants are more likely to have genetic predispositions correlated with lower EA and worse health than migrants. Distinguishing our findings from previous work, we show that migrants from “rich” to “poor” places have higher measures of PGI for EA than do those who remain (non-migrants) in rich places—showing that genetics selects whether people migrate, even those migrating to “deprived” areas.

## Results

### Re-classification of Migration Pattern

Using data from the UK Biobank (UKB), we reassessed AA’s work with a revised migration classification based on the “type” of place of residence at birth compared with at the time of the survey; places were either “deprived” (defined as a coal-mining place) or not. Following prior research, we first classified UKB respondents into four migration groups: (1) moved away from coal mining place; (2) stayed out of coal mining place; (3) moved to coal mining place; and (4) stayed in coal mining place. We note that the second and fourth (“stay”) categories combine respondents who did not move and those who moved but remained in the same “type” of place (e.g., moved between two coal mining places). Thus, the four-type classification cannot separate the analyses of genetic differences in whether UKB respondents move and genetics differences in where they move (or stay).

Our analysis aims to test if genetics differ in whether respondents migrate, or where they migrate, or both. To do so, we updated the four-category migration measure to a six-category migration measure: (A) moved from coal mining to non-coal mining place; (B) moved between two non-coal mining places; (C) stayed in the same non-coal mining place; (D) moved from non-coal mining to coal mining place; (E) moved between two coal mining place; and (F) stayed in the same coal mining place. Note that groups B and E were classified as stayers (i.e., non-migrants) in earlier work. A summary of migration classification and the sample size of each migration group are available in Table 1.

**Table 1:**
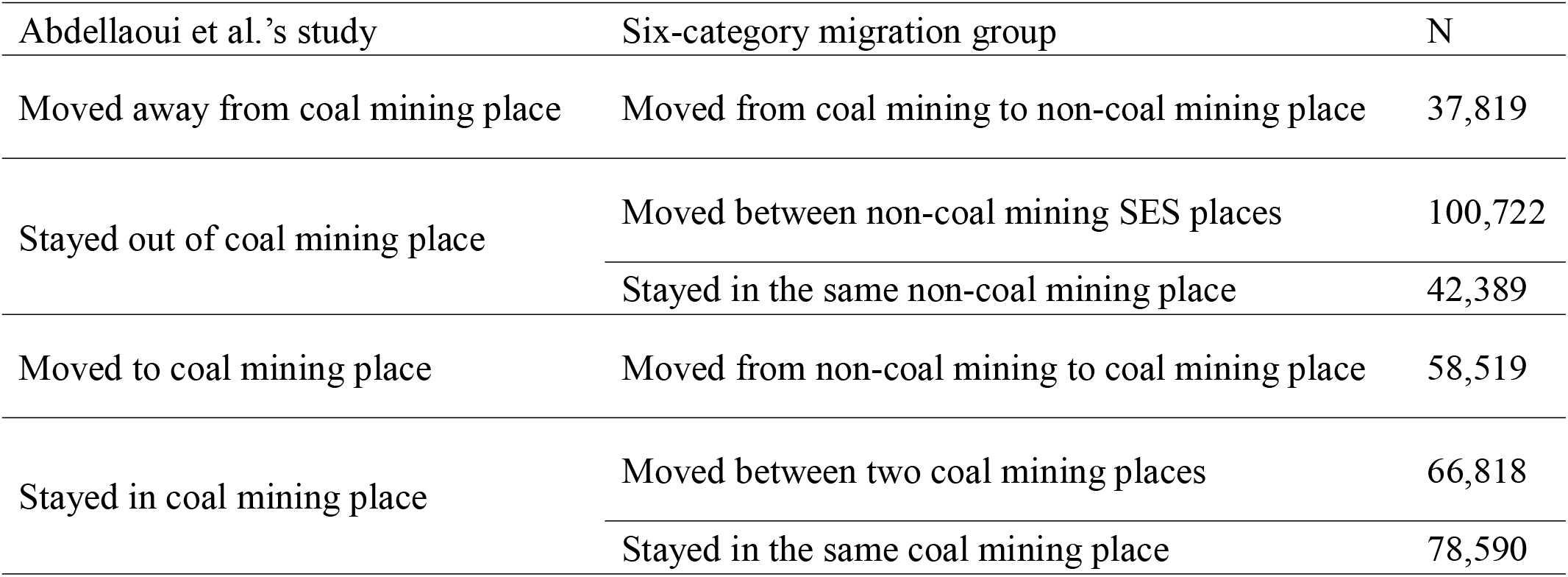
Comparison of classifications of internal migration patterns in the UK Biobank.

### Migration and EA PGI

Predicted EA PGI by the six-category migration group are presented in Figure 1 (estimated regression coefficients are available in Supplementary Table 1). Results of the six-category migration group show differences between migrants and non-migrants in both coal mining and non-coal mining places. For example, among those who are living in a non-coal mining place, non-migrants’ EA PGI (group C) is substantially lower than migrants’ EA PGI (groups A and B). Similar migrant and non-migrant differences can be seen for those who are living in a coal mining place (groups D–E vs. group F).

**Figure 1:**
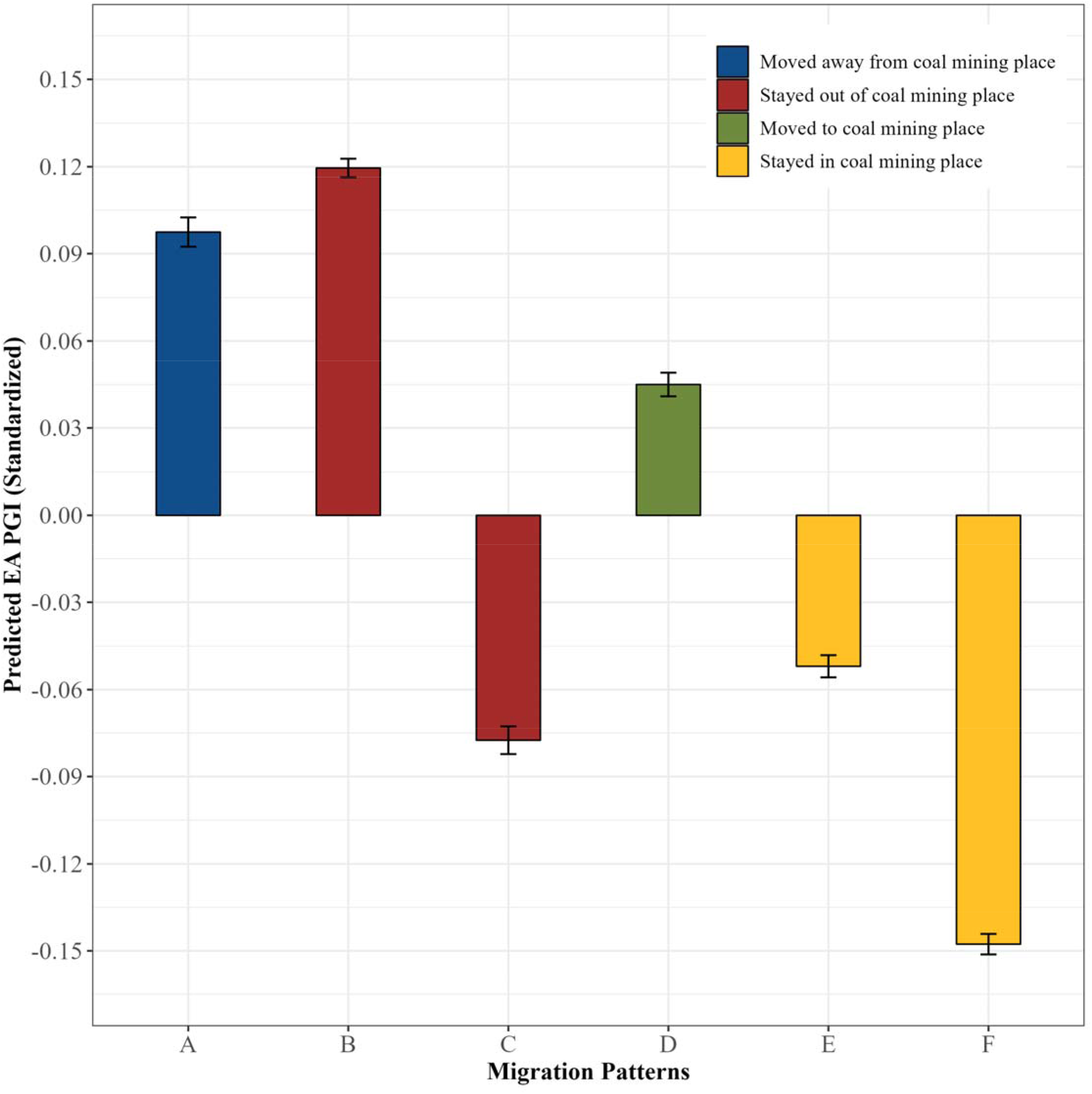
Migration patterns and educational attainment polygenic index. Note: Bars and error bars represent estimated mean polygenic indices and their robust standard errors. First 20 principal components are controlled. Capital letters indicate the six-category migration groups: (A) moved from coal mining to non-coal mining place; (B) moved between two non-coal mining places; (C) stayed in the same non-coal mining places; (D) moved from non-coal mining to coal mining place; (E) moved between two coal mining place; and (F) stayed in the same coal mining place.

We can also observe a clear difference between migrants and non-migrants by place of birth. Specifically, among those who were born in a non-coal mining place, migrants (groups B and D) carry a significantly and substantially higher EA PGI than non-migrants (group C). Similarly, among those who were born in a coal mining place, migrants’ EA PGI (groups A and E) is significantly higher than non-migrants’ EA PGI (group F). Furthermore, those who stayed in the same non-coal mining place (group C) have a significantly lower EA PGI than those who moved between two coal mining places (group E). Overall, these results suggest that, regardless of “type” of birthplace and place of residence at the time of survey, migrants have genetic variants correlated with better EA than non-migrants.

Results of the six-category migration group also demonstrate within-migrants differences. Among migrants from a coal mining place, those who moved to a non-coal mining place (group A) have a significantly higher EA PGI than their counterparts who moved to another coal mining place (group E). Likewise, there is also a significant difference in EA PGI by the “type” of place of residence at the time of survey among migrants from a non-coal mining place (groups B and D). We can also see differences by where migrants came from (groups A and B, and groups D and E). The size of difference is not as large as that of the differences in EA PGI by where migrants moved to, but migrants born in a non-coal mining place have higher EA PGI than migrants born in a coal mining place.

We then turned to the reassessment of the association between EA PGI and migration patterns over time. Figure 2 Panel A shows a decline in mean EA PGI over time, unlike an increase in years of schooling in Figure 2 Panel D. These findings are consistent with AA, suggesting the presence of the negative association between fertility rate and EA PGI (Kong et al., 2017) and survival selection. Figure 2 Panel B illustrates that the significant differences in EA PGI between migrants and non-migrants persist across birth cohorts. Further, non-migrants contribute more to the decline in mean EA PGI over time than migrants. On average, among migrants, each year increase in birth year is associated with 0.00134 standard deviations decrease in EA PGI, whereas the corresponding one-year decrease in EA PGI for non-migrants is 0.00276 standard deviations. This slope difference between migrants and non-migrants is statistically significant at the 0.1% level. The steeper slope for non-migrants implies the growing difference in EA PGI between migrants and non-migrants in more recent birth cohorts.

**Figure 2:**
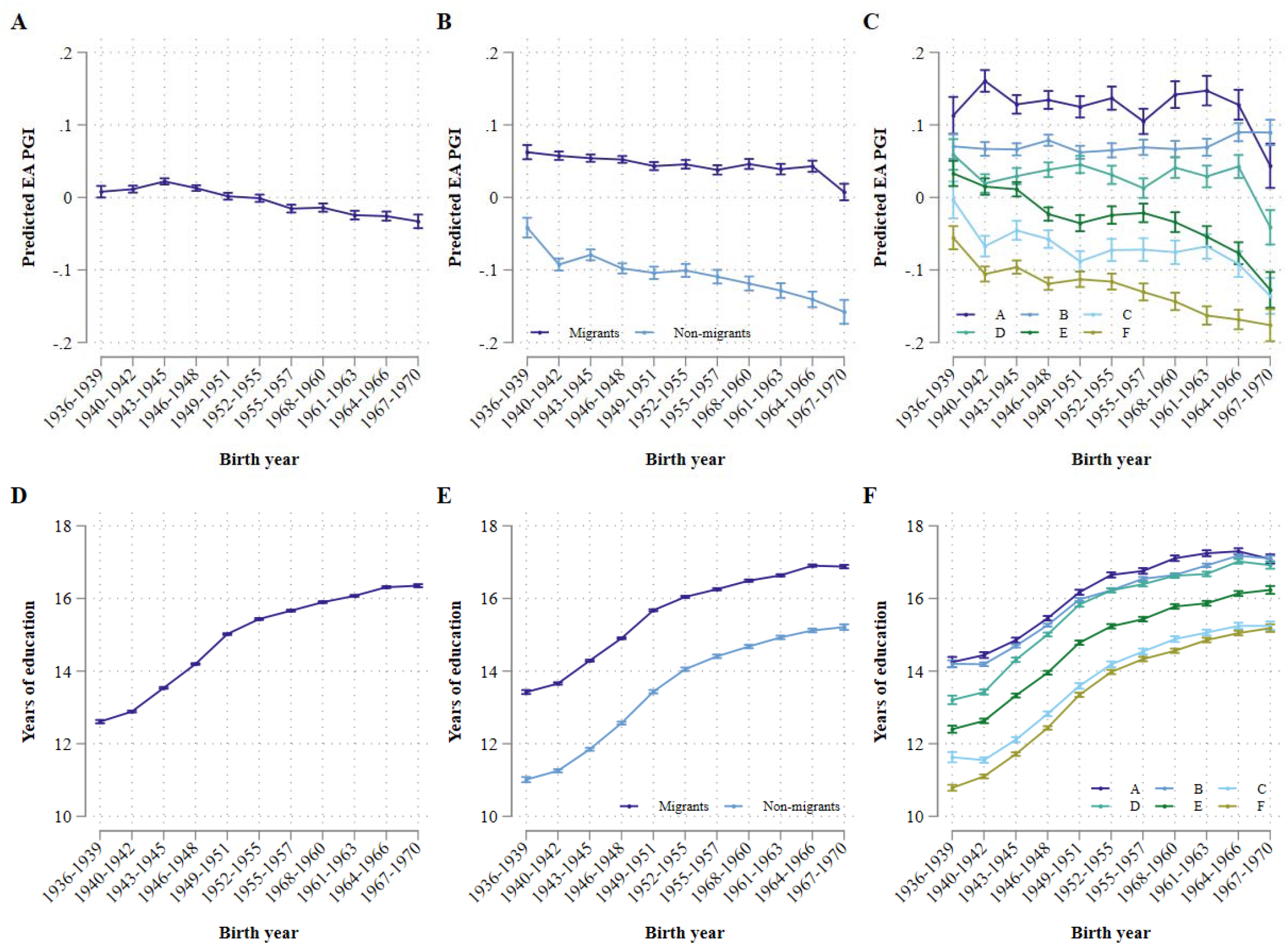
Estimated mean educational polygenic index and years of schooling, by birth year. Note: Dots and error bars represent estimated mean and their robust standard errors. First 20 principal components are controlled for educational attainment polygenic indices. Capital letters indicate the six-category migration groups: (A) moved from coal mining to non-coal mining place; (B) moved between two non-coal mining places; (C) stayed in the same non-coal mining places; (D) moved from non-coal mining to coal mining place; (E) moved between two coal mining place; and (F) stayed in the same coal mining place. Years of education are based on International Standard Classification of Education (Lee et al., 2018).

Figure 2 Panel C presents the association between EA PGI and birth year by the six-category migration group. Consistent with the results in AA, we observed steeper decreases in EA PGI among those were born and living in a coal mining place (groups E and F); however, there also exists a decline in EA PGI among non-migrants in a non-coal mining place (group C). For those who were born and stayed in the same non-coal mining place, the association between birth year and EA PGI is statistically significant at the 0.1% level (β = -0.0022). By contrast, the corresponding regression coefficients for the other groups (groups A, B, and D) are relatively small (ranging from -0.0006 to -0.0011) and not statistically significant at the conventional 5% significance level (p = 0.0844 for group A, p = 0.0982 for group B, and p = 0.247 for group D). It is important to note that EA PGI of those who stayed in the same non-coal mining place (group C) is consistently lower than migrants who moved to a coal mining place (group D). Overall, these results suggest that non-coal mining places do not send those with lowest EA PGI to coal mining places over time, rather they keep those with even lower EA PGI in non-coal mining places in more recent birth cohorts.

### Migration and PGI for Health-Related Traits

Because migration is also a selective process in health (Jasso et al., 2004; Palloni & Arias, 2004; Palloni & Morenoff, 2006), we conducted parallel analyses with 12 health-related PGIs. Figure 3 summarizes the relationships between migration and 12 health-related PGIs (estimated regression coefficients are available in Supplementary Tables 1). Although less clear, results are generally consistent with the findings with EA PGI. Specifically, non-migrants who stayed out of coal mining places (group C) have PGIs correlated with a lower level of cognitive function, higher body mass index (BMI) and waist-to-hip ratio, higher risks of attention-deficit/hyperactivity disorder (ADHD) and coronary artery disease (CAD), shorter height, higher likelihood of ever smoked, and more frequent cigarette smoking than migrants moved to a coal mining place (group D) at the 5% level. Further, non-migrants in coal mining places (group F) have genetic variants correlated with lowest level of health relative to any migrant groups. Importantly, genetic risks of health issues are generally higher for migrants than for non-migrants; however, the genetic risk of bipolar disorder is an exception. Non-migrants have a lower genetic risk of bipolar disorder than migrants.

**Figure 3:**
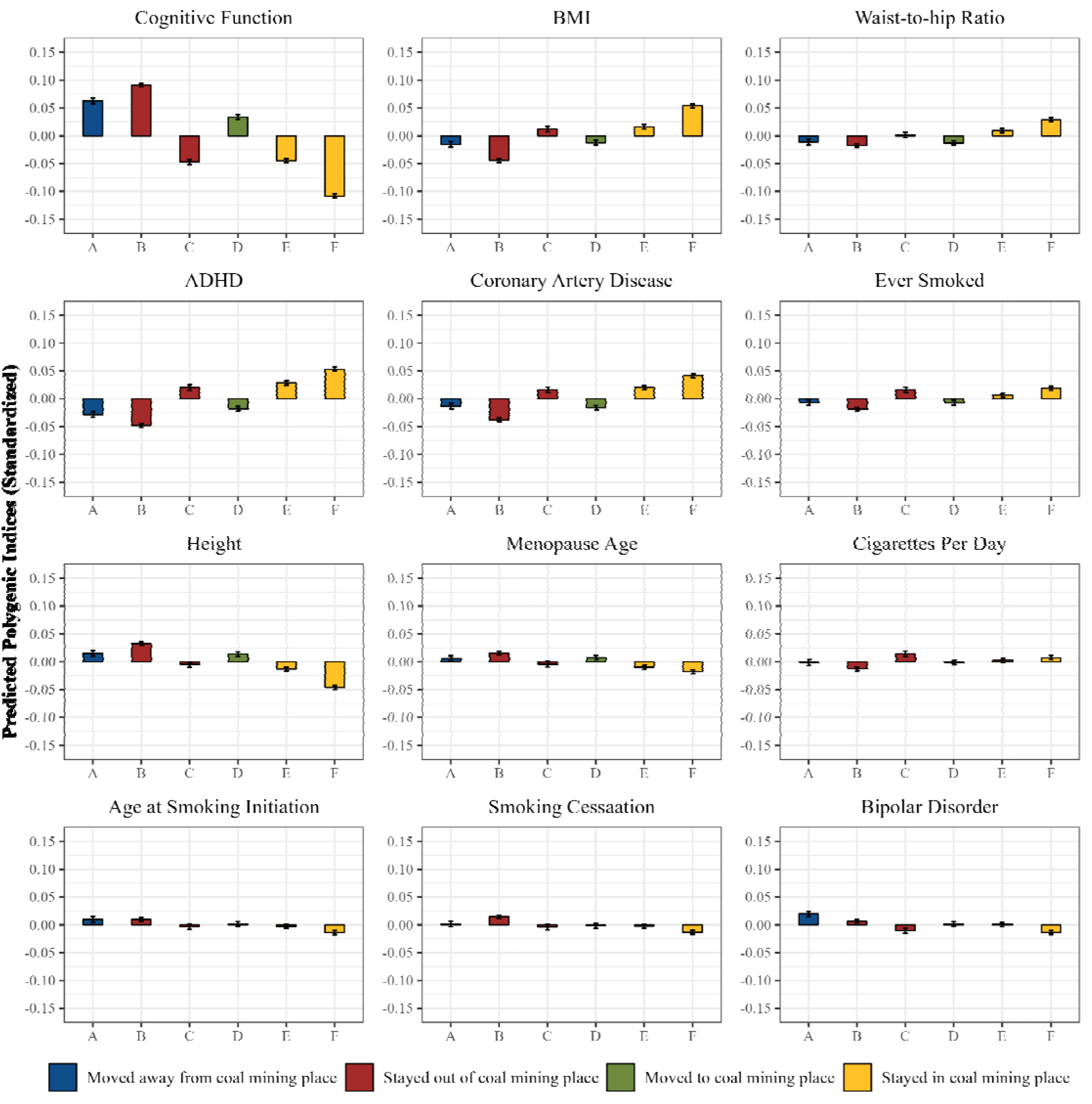
Migration patterns and health-related polygenic indices. Note: Bars and error bars represent estimated mean polygenic indices and their robust standard errors. First 20 principal components are controlled. Capital letters indicate the six-category migration groups: (A) moved from coal mining to non-coal mining place; (B) moved between two non-coal mining places; (C) stayed in the same non-coal mining places; (D) moved from non-coal mining to coal mining place; (E) moved between two coal mining place; and (F) stayed in the same coal mining place.

We also observe within-migrant differences. Among migrants born in a coal mining place, those who moved to a non-coal mining place (group A) have significantly higher cognitive function, height, and age of menopause PGIs and lower BMI, waist-to-hip ratio, ADHD, CAD, and ever smoked PGIs than those who moved between two coal mining places (group E). Similarly, those who moved between two non-coal mining places (group B) have significantly higher cognition, height, and smoking cessation PGIs and lower PGIs correlated with BMI, ADHD, CAD, ever smoked, and number of cigarettes per day relative to those who moved from non-coal mining to coal mining place (group D). Further, among migrants living in a non-coal mining place, those born in a non-coal mining place (group B) generally have genetic variants correlated with better health than their counterparts born in a coal mining place (group A). Similar differences can be seen among migrants living in a coal mining place (groups D and E).

## Discussion

Geographic mobility is one of the mechanisms shaping the geographic distribution of genetic predispositions. AA has demonstrated the role of geographic mobility in the geographic distribution of genetic predisposition; however, their migration measure does not clearly distinguish migrants from non-migrants. The goal of this study is to deepen our understanding of the relationship between migration, social stratification, and genetic predispositions through a reassessment of AA’s work with an updated migration measure distinguishing migrants from non-migrants.

First, we assessed how EA PGI differs by where people moved. Consistent with the findings in AA’s work, we found that those with high EA PGI tend to leave economically deprived areas. Further, our results also showed that, among migrants born in an economically prosperous area, those with low EA PGIs more likely move to a deprived area. These findings conditionally support the two implications provided by AA. That is, *among migrants*, own genetic variants are more similar to neighbors’ genetic variants than those living far away, and migration contributes to geographic clustering of genetic variants.

We also assessed how EA PGI differs by whether people moved. We found that those with lowest EA PGIs do not move out but stay in their birthplaces, regardless of SES of birthplace. Given the strong association between EA and health (Barrow & Malamud, 2015; Elo, 2009; Hout, 2012), these findings are consistent with the healthy migrant literature (Jasso et al., 2004; Palloni & Arias, 2004; Palloni & Morenoff, 2006), suggesting that skilled and healthy individuals are more likely to migrate than unskilled and unhealthy counterparts. The low EA PGI among non-migrants provide some caveats to the two key interpretations of AA’s work. First, neighbors’ PGI is not always similar to own PGI than the PGI of people living far apart. Specifically, EA PGI for non-migrants in less economically deprived areas is not as high as EA PGI for migrants in less economically deprived areas, but as low as EA PGI for those in economically deprived areas. Combined with the findings that migrants living in non-coal mining places have higher EA PGI than migrants in coal mining places, our results demonstrated that migrants move to places where their own genetic predispositions are similar to neighbors’, but non-migrants stay in the same place even though their neighbors’ genetic predispositions are substantially different from their own genetic traits. Second, geographic clustering of PGI through migration is in line with recent SES-driven migration (Coulter & Scott, 2015; Rodden, 2010) only among migrants leaving coal mining places when we took non-migrants into consideration. Because non-coal mining places do not send a group of individuals whose PGI are correlated with lowest EA, migration moving to coal mining places is not consistent with the finding that the durable and affordable housing and public transportation networks in coal mining places attract poor, low-skilled migrants (Rodden, 2010). Hence, migration contributes to geographic clustering of those with low EA PGI in economically deprived areas less than AA reported.

We also investigated relationships between migration patterns and health-related PGIs. Our results for PGIs correlated with health outcomes also showed that migrants moved out from a coal mining place have PGIs correlated with better health, but non-coal mining places do not send a group of individuals with genetic variants correlated with worst health to coal mining places. These results are generally consistent with our findings for EA PGI. Whereas migration out from a coal mining place facilitates the geographic clustering of PGIs correlated with better health in economically prosperous areas, geographic clustering of PGIs correlated with worse health in economically deprived areas through migration is less supported. Importantly, as AA reported, we confirmed that less deprived areas attract migrants with a high bipolar disorder PGI. The high genetic risk of bipolar disorder among those who moved out from a coal mining place is not consistent with results of the other health-related PGIs. One possible interpretation of this unanticipated result is that skilled individuals are more likely to suffer from bipolar disorder. This implies that less deprived areas attract skilled individuals who also have high genetic risk of bipolar disorder. This interpretation is consistent with findings in epidemiology which have demonstrated a higher risk of bipolar disorder among skilled individuals (MacCabe et al., 2010; Tiihonen et al., 2005).

This study is not without limitations. First, the UKB is not a nationally representative survey. The UKB recruited only those living reasonably close to assessment centers, and oversampled well-educated and healthy UK residents (Fry et al., 2017; Munafò et al., 2018). The UKB’s unique sampling strategy may induce biased estimates; therefore, our findings may not be representative of the relationships between migration and genetic variants in the UK. Additionally, the unequal probability to participate in the UKB may induce spurious relationships between migration and genetics if both migration and genetics affect the probability to participate in the UKB. Nevertheless, we are uncertain how the unique sampling strategy affects the compositions of migration groups and our findings, as AA also acknowledged. Subsequent research replicating our findings with a nationally representative survey is, therefore, an important extension of this study.

Second, non-European ancestry respondents were excluded from our analytical sample to increase ancestral homogeneity. The exclusion of non-European ancestry individuals prevents us from generalizing our findings because we cannot infer the relationship between migration and genetic predispositions to non-European ancestral groups. This is an important limitation given that unequal advancement of genetic research by ancestral groups may exacerbate existing disparities between Europeans and other ancestral groups (Martin et al., 2019).

Finally, our migration measures relied on place of birth and current residence at the time of the survey, but this snapshot precludes us from capturing migration history. Respondents who had migrated to pursue college education in a non-coal mining place but returned to their coal mining birthplace to work may carry different genetic variants from those who stayed in their coal mining birthplace over the life course. This is a well-known measurement error in migration research (Molloy et al., 2011), and lifetime migration history information would allow us to evaluate the relationship between geographic mobility and genetic variants in a greater detail; however, such data do not exist.

Despite these limitations, the current study advances our understandings of associations between migration and genetic predispositions. Our findings showed that economically deprived areas send individuals with high genetic propensities to achieve higher socioeconomic status to less economically deprived areas, whereas economically prosperous areas do not send individuals with genetic variants correlated with lowest socioeconomic status and worst health. These findings imply that a cycle of cumulative advantage of less economically deprived areas is only partially supported. Nevertheless, this does not indicate the absence of exacerbation of social inequalities at the genetic level between places through geographic mobility because people with genetic variants correlated with higher socioeconomic status are leaving economically deprived areas. This highlights the need of continued scholarly efforts to investigate the role of migration in geographic inequality of genetic variants.

## Methods

### Sample

We used data from the UKB, in conjunction with data on geographic boundaries of coal fields (https://data.gov.uk/dataset/coal-mining-reporting-area) and local authorities for 2011 Census in the UK (http://infuse.ukdataservice.ac.uk/help/definitions/2011geographies/index.html). The UKB is a large-scale, population-based, longitudinal biobank study of over 500,000 people and collected baseline data from 2006 to 2010. The UKB recruited the baseline sample through an invitation letter to 40–69 years old individuals who registered with a National Health Service General Practitioner and lived reasonably close to one of the 22 catchment areas where UKB assessment centers were located. Of completed respondents in the UKB (n = 502,505), we excluded individuals of non-European ancestry (n = 93,549) to increase ancestral homogeneity of our sample. Additionally, we also excluded those without place of birth coordinates (n = 23,421) and lacking genetic data (n = 678) which left us an analytical sample of 384,857 respondents.

### Variables

#### Genetic Measures

We used polygenic indices as a genetic marker to re-evaluate how migration patterns are associated with genetics. A PGI represents cumulative effects of independent genomic loci of small effect. We used results of GWAS summary statistics in published papers to construct 13 specific PGIs: EA (Lee et al., 2018), cognitive function (Rietveld et al., 2014), BMI (Locke et al., 2015), waist-hip ratio (Shungin et al., 2015), ADHD (Demontis et al., 2019), CAD (Nikpay et al., 2015), ever smoked (The Tobacco and Genetics Consortium, 2010), height (Wood et al., 2014), age at menopause (Day et al., 2015), number of cigarette per day, age of smoking initiation, smoking cessation (The Tobacco and Genetics Consortium, 2010), and bipolar disorder (Stahl et al., 2019). To avoid overfitting, we ensured that UKB samples were excluded from GWAS summary data used for PGI training. We clumped single-nucleotide polymorphisms (SNPs) using Phase III European samples from 1,000 Genomes Project as the linkage disequilibrium (LD) reference. The LD window size of 1 Megabase (Mb) and a pairwise r2 threshold of 0.1 were used. No p-value thresholding was applied for variant selection. The PGIs were calculated using PRSice-2 (Choi & O’Reilly, 2019) and were standardized to a mean of 0 and a variance of 1 in downstream analyses.

#### Migration Patterns

We used coordinate variables of place of birth and current residence at the time of initial survey interview (2006–2010) to construct a migration pattern variable, and coordinate information in a follow-up survey for a small portion of respondents (n = 11) who did not provide them in an initial survey but provided in a follow-up survey. Using the coordinate and geographical boundary data, we classified UKB respondents into the six-category migration group: (A) moved from low to high SES place; (B) moved between two high SES places; (C) stayed in the same high SES place; (D) moved from high to low SES place; (E) moved between two low SES places; and (F) stayed in the same low SES place. In doing so, we considered two geographical dimensions. First, we identified whether respondents’ birthplace and place of residence are low SES (i.e., coal mining) or high SES (i.e., non-coal mining) places. We achieved this by using geographical boundary data for coal mining places. We then classified respondents as non-migrants if they lived in the same local authorities as where they were born; otherwise, they were classified as migrants. We employed the local authorities used in the 2011 Census as a geographical unit because we have publicly available geographical boundary data. In the 2011 Census, there were 404 local authorities and the average land area of local authority districts in 2015 was approximately 240 square miles. This is close to the average land area of counties in Rhode Island (207 square miles) and Virginia (295 square miles) which are second and third smallest average land area within a state, just following District of Columbia.

### Analytical Strategy

To test the differences in genetic characteristics by migration groups, we used an ordinary least squares regression. The regression equation can be expressed as follows:

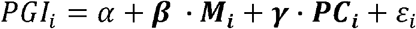

where *PGI*_*i*_ is a polygenic index; ***M***_*i*_ is a dummy-coded migration group assignment; and ***PC***_*i*_ is a vector of the first 20 genetic principal components which account for population structure-related confounding effects (Price et al., 2006). We then added a birth year variable and an interaction between birth year and migration patterns to assess longitudinal changes in the association between educational attainment polygenic index and migration patterns.

## Acknowledgement

The authors gratefully acknowledge use of the facilities of the Center for Demography of Health and Aging (P30 AG016266) and the Center for Demography and Ecology (P2C HD067873) at the University of Wisconsin–Madison. We thank members of the Social Genomics Working Group at University of Wisconsin for helpful comments. We also thank Abdel Abdellaoui and colleagues for providing programming code and geographical boundary data to reassess their work. Earlier version of this paper was presented in the Population Association of America 2021 Annual Meeting. Results from this research were also presented earlier at the National Institute on Aging supported 2021 Integrating Genetics and Social Sciences Conference (R13-AG062366). This research has been conducted using the UK Biobank Resource under Application 57284.

## Data Availability

This research has been conducted using the UK Biobank resource. UK Biobank data can be accessed on request with an approval by the UK Biobank committee. The regional measures are publicly available through the URL in the Methods section.

## Code Availability

Stata and R code used for statistical analyses are available upon request.

## Notes

### Competing Interest Statement

The authors have declared no competing interest.

