## Supplementary Table 1 for "The Big (Genetic) Sort? Reassessing Migration Patterns and Their Genetic Imprint in the UK"

**Supplementary Materials**

### Supplementary Table 1: Regression coefficients of six migration groups for polygenic indices.

| VARIABLES | Educational attainment | Cognitive function | BMI | Waist-to-hip ratio |
| --- | --- | --- | --- | --- |
| Migration groups (ref: Moved from low to high SES places) |  |  |  |  |
| Moved between two high SES places | 0.022*** | 0.028*** | -0.029*** | -0.006 |
|  | (0.006) | (0.006) | (0.006) | (0.006) |
| Stayed in the same high SES place | -0.052*** | -0.029*** | 0.003 | -0.002 |
|  | (0.007) | (0.007) | (0.007) | (0.006) |
| Moved from high to low SES places | -0.149*** | -0.108*** | 0.032*** | 0.020** |
|  | (0.006) | (0.006) | (0.006) | (0.006) |
| Moved between two low SES places | -0.175*** | -0.110*** | 0.028*** | 0.013† |
|  | (0.007) | (0.007) | (0.007) | (0.007) |
| Stayed in the same low SES place | -0.245*** | -0.171*** | 0.069*** | 0.040*** |
|  | (0.006) | (0.006) | (0.006) | (0.006) |
| N | 384,857 | 384,857 | 384,857 | 384,857 |

Note: First 20 genetic principal components are controlled. Additional controls are not shown. Robust standard errors are in parentheses. *** p<0.001, ** p<0.01, * p<0.05 (two-tailed).

(Cont. Supplementary Table 1)

| VARIABLES | ADHD | CAD | Ever smoked | Height |
| --- | --- | --- | --- | --- |
| Migration groups (ref: Moved from low to high SES places) |  |  |  |  |
| Moved between two high SES places | -0.020** | -0.024*** | -0.012* | 0.018** |
|  | (0.006) | (0.006) | (0.006) | (0.006) |
| Stayed in the same high SES place | 0.010 | -0.002 | -0.000 | -0.002 |
|  | (0.007) | (0.007) | (0.006) | (0.006) |
| Moved from high to low SES places | 0.057*** | 0.034*** | 0.013* | -0.029*** |
|  | (0.006) | (0.006) | (0.006) | (0.006) |
| Moved between two low SES places | 0.048*** | 0.030*** | 0.023*** | -0.020** |
|  | (0.007) | (0.007) | (0.007) | (0.007) |
| Stayed in the same low SES place | 0.081*** | 0.055*** | 0.026*** | -0.061*** |
|  | (0.006) | (0.006) | (0.006) | (0.006) |
| N | 384,857 | 384,857 | 384,857 | 384,857 |

Note: First 20 genetic principal components are controlled. Additional controls are not shown. Robust standard errors are in parentheses. *** p<0.001, ** p<0.01, * p<0.05 (two-tailed).

(Cont. Supplementary Table 1)

| VARIABLES | Menopause age | Cigarettes per day | Age at smoking initiation | Smoking cessation |
| --- | --- | --- | --- | --- |
| Migration groups (ref: Moved from low to high SES places) |  |  |  |  |
| Moved between two high SES places | 0.010† | -0.011† | -0.000 | 0.013* |
|  | (0.006) | (0.006) | (0.006) | (0.006) |
| Stayed in the same high SES place | 0.002 | 0.000 | -0.008 | -0.003 |
|  | (0.007) | (0.007) | (0.007) | (0.007) |
| Moved from high to low SES places | -0.015* | 0.004 | -0.012† | -0.004 |
|  | (0.006) | (0.006) | (0.006) | (0.006) |
| Moved between two low SES places | -0.010 | 0.016* | -0.013† | -0.006 |
|  | (0.007) | (0.007) | (0.007) | (0.007) |
| Stayed in the same low SES place | -0.023*** | 0.009 | -0.023*** | -0.015* |
|  | (0.006) | (0.006) | (0.006) | (0.006) |
| N | 384,857 | 384,857 | 384,857 | 384,857 |

Note: First 20 genetic principal components are controlled. Additional controls are not shown. Robust standard errors are in parentheses. *** p<0.001, ** p<0.01, * p<0.05 (two-tailed).

(Cont. Supplementary Table 1)

| VARIABLES | Bipolar disorder |
| --- | --- |
| Migration groups (ref: Moved from low to high SES places) |  |
| Moved between two high SES places | -0.013* |
|  | (0.006) |
| Stayed in the same high SES place | -0.018** |
|  | (0.006) |
| Moved from high to low SES places | -0.019** |
|  | (0.006) |
| Moved between two low SES places | -0.030*** |
|  | (0.007) |
| Stayed in the same low SES place | -0.033*** |
|  | (0.006) |
| N | 384,857 |

Note: First 20 genetic principal components are controlled. Additional controls are not shown. Robust standard errors are in parentheses. *** p<0.001, ** p<0.01, * p<0.05.
